# Alginate hydrogels loaded with pyruvate dehydrogenase enzyme: a novel wound dressing to disassemble *Pseudomonas aeruginosa* biofilm

**DOI:** 10.1101/2025.06.26.661844

**Authors:** Omid Sedighi, Nicole Roode, Tucker Johnsen, Maisie Moses, Karin Sauer, Amber L. Doiron

## Abstract

Topical wounds require effective dressings to manage wound exudate, maintain moisture, and avoid infection. Alginate hydrogels are widely used clinically due to their biocompatibility, high absorbency, and ability to maintain a moist wound environment during healing; however, alginate can also promote bacterial biofilm formation and prolong healing. Depleting pyruvate using the enzyme pyruvate dehydrogenase (PDH) has been shown in both *in vitro* and *in vivo* studies to effectively disrupt biofilm structure. This study entraps PDH in alginate hydrogels to disperse/disassemble biofilms formed by a prevalent wound pathogen, *Pseudomonas aeruginosa*. PDH depletes pyruvate and alters the biofilm metabolism, resulting in the prevention and dispersion of biofilms. Alginate hydrogels exhibited compressive moduli ranging from 15 to 80 kPa, with the water content of all swollen hydrogels exceeding 90%, and demonstrated structural stability in simulated wound exudate. PDH loading of 0.85 U in 65 µl of alginate achieved maximum observed enzymatic activity, 60% of which was retained after six days. Notably, PDH-loaded hydrogels significantly reduced biofilm biomass by 40.94% ± 6.07 compared to media alone and by 80.05% ± 6.37 compared to the Dimora® commercial alginate dressing. Furthermore, PDH-entrapped alginate hydrogels significantly reduce biofilm biomass without affecting the viability of human dermal fibroblasts, highlighting the commercial potential of the material. The successful integration of a biofilm metabolism-altering enzyme within wound dressings presents a promising avenue for improving chronic wound management and reducing the burden of biofilm-associated infections without the need for the discovery of new antibiotic drugs.

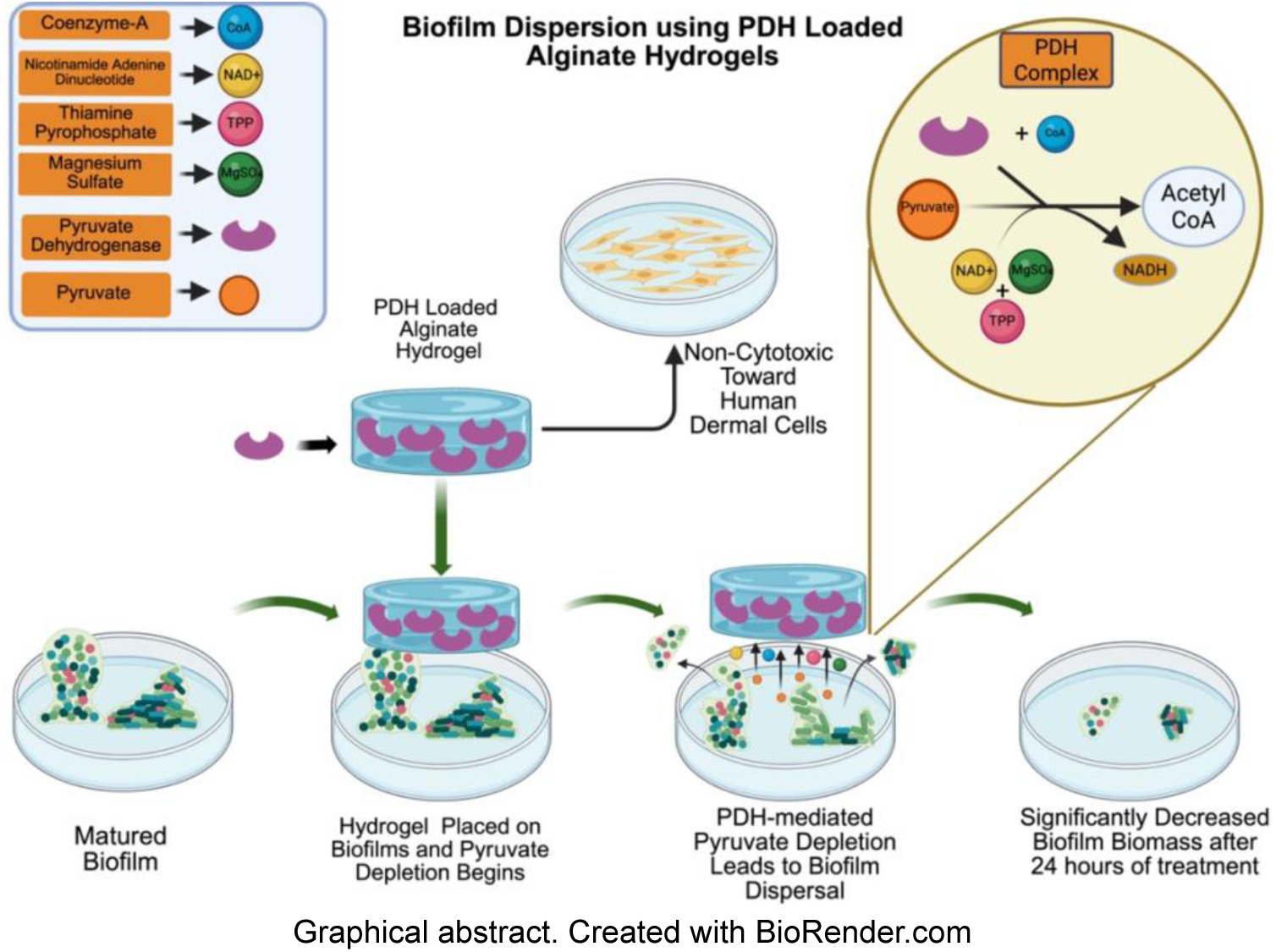

## Introduction

Skin is a vital barrier against external pathogens, and upon injury, a complex and highly regulated healing process begins [1, 2]. Wound dressings play a critical role in maintaining an optimal environment for healing and protecting the wound from infection. Due to their excellent swelling capacity and ability to maintain a moist environment, hydrogels are one of the primary material types used in wound dressings. An ideal wound dressing must meet several key criteria: sufficient mechanical integrity to withstand patient movement, conformability to the wound, high absorbency to manage wound exudate and retain absorbed fluids, minimal degradation during usage to ensure prolonged coverage, and limited potential for infection [3].

Alginate is a commonly used biopolymer in FDA-approved wound dressings that is biocompatible, non-immunogenic, highly absorbent, and maintains a physiologically moist environment to promote wound healing. Despite its widespread clinical use, alginate can encourage the development of bacterial communities known as biofilms. Biofilms are surface-associated bacterial communities that are reticent to treatment and are a leading cause of non-healing chronic wounds [4, 5]. The antibiofilm properties of several commercially available wound dressings, including the calcium alginate dressing Algisite®-M, have been evaluated in comparison to no dressing and wet gauze [6]. With an initial biofilm biomass of 7.076 ± 0.436 log CFU/ml, after 24 hours of treatment, the biofilm biomass increases to 9.051 ± 0.204 log CFU/ml (∼94-fold increase) with Algisite®-M, 6.075 ± 0.324 log CFU/ml (10-fold decrease) with no dressing, and 7.827 ± 0.385 log CFU/ml (∼5.6-fold increase) with wet gauze. Given the widespread use of alginate-based wound dressings in clinical settings, this enhanced biofilm growth is concerning, as it may negatively impact wound healing and patient recovery. To address this issue, a common strategy is to incorporate silver nanoparticles into alginate dressings. In the same study, Acticoat® Absorbent—a commercially available silver-loaded alginate dressing designed to release 70–100 ppm of nanocrystalline silver ions over 7 days—causes biofilm biomass to reach 6.206 ± 0.144 log CFU/l (∼7-fold decrease) after 24 hours. Thus, while the addition of silver reduced biofilm formation relative to plain alginate, the biofilm biomass still increased over time and remained higher than the no-dressing control. In a separate study on *E. coli, S. aureus, P. aeruginosa, Streptococcus gordonii, Streptococcus mitis, Streptococcus mutans, Enterococcus faecalis*, and *Candida albicans*, the incorporation of silver nanoparticles results in a 44-61% reduction in biofilm mass after 24 hours of treatment; however, this reduction was fund to be highly bacterial strain-specific [7].

Although silver has demonstrated significant antimicrobial efficacy both *in vitro* and *in vivo*, when incorporated into dressings such as alginate, it does not exhibit strong antibiofilm activity. Moreover, silver can be cytotoxic to host cells at high concentrations, making controlled release essential [8]. Other strategies such as the addition of lactoferrin (LTF) [9], incorporation of silica [10], and stabilization with tannic acid [11] have been explored to enhance silver stability and its antibiofilm properties, but still remain under investigation. LTF, an iron cleaving protein, which can enhance silver’s antibiofilm properties by depriving iron. Moreover, LTF can bind with lipopolysaccharide, leading to the disruption of bacterial biofilms and prevention of re-colonization [12]. Incorporation of silica prevents AgNPs from aggregation and controls its release allowing for prolonged antimicrobial effects [10]. Stabilization of AgNPs in tannic acid also improves their long-term usage by preventing aggregation [11]. In addition to AuNP, the incorporation of nitric oxide (NO) donors into alginate dressings can enhance antibacterial properties [13, 14], yet there are limitations, such as the short half-life of NO itself and potential off-target effects [15, 16]. The addition of antimicrobial peptides (AMPs) such as human β-defensin 2 (hBD-2) and PP4-3.1 [17], D-Bac8c^2,5 Leu^, [18], and Chol-37 (F34-R) [19] to alginate has also been studied. Yet, AMPs face several limitations that hinder their clinical and commercial development, including potential toxicity, reduced activity in the presence of wound serum, susceptibility to proteases, and challenges in delivery and bioavailability [20]. Adding bacteriophages to alginate hydrogels can also add antibiofilm properties to the hydrogels [21, 22]. But, this method is limited to particular bacterial species or even single strains [23]. In this work, we developed a metabolism-altering wound dressing using the enzyme pyruvate dehydrogenase (PDH), which can potentially address the above-mentioned shortcomings of alginate wound dressings.

Our previous work supports the use of pyruvate depletion as a therapeutic approach to alter biofilm metabolism and induce dispersion. Dispersion is the last phase of biofilm formation during which more than 80% of the biofilm’s biomass can be shed, and the dispersed bacteria become as susceptible to antibiotics as those in the planktonic state [24]. Pyruvate depletion, achieved through the enzyme PDH, coincides with a significant reduction in the biomass of *P. aeruginosa* biofilms [25]. Similar biomass reductions are observed when pyruvate depletion is induced in *Staphylococcus aureus* (*S. aureus*), causing 40% of biofilms to disperse when exposed to PDH compared to 8% in untreated colonies [26]. PDH has previously been encapsulated in poly(lactic-co-glycolic) nanoparticles (PLGA NPs) as a means of stabilizing enzyme activity during storage in a drug delivery platform [27]. In addition to this stabilization, PLGA NPs-encapsulated PDH disperse *P. aeruginosa* biofilms at levels comparable to those of unencapsulated PDH. This previous work motivates the current study, which focuses on mitigating the adverse outcomes of alginate wound dressings attributed to biofilms.

In this work, the mechanical, chemical, and physical characterization of PDH-loaded alginate hydrogels is summarized, along with biofilm growth assays, which compare the synthesized gels to clinically marketed dressings. Finally, biocompatibility with *in vitro* human dermal fibroblasts has been established, demonstrating the potential for this technology to disaggregate biofilms in the presence of these alginate wound dressings.

## Materials and Methods

### Materials

Calcium chloride (CaCl_2_), sodium pyruvate, sodium chloride (NaCl), sodium alginate (Na-Alg), pyruvate dehydrogenase from porcine heart (PDH), coenzyme A sodium salt hydrate (CoA), thiamine pyrophosphate (TPP), magnesium sulfate (MgSO_4_), bovine serum albumin (BSA), 3-(N-Morpholino) propanesulfonic acid sodium salt (MOPS), and β-nicotinamide adenine dinucleotide hydrate (β-NADH) were purchased from Sigma-Aldrich (St. Louis, MO, USA). Crystal violet (CV, certified biological stain), BD Difco™ LB Broth, and tris(hydroxymethyl)aminomethane were purchased from Fisher Scientific (Waltham, MA, USA). The Micro BCA™ Protein Assay Kit was purchased from Thermo Fisher Scientific (Waltham, MA, USA). *P. aeruginosa* (PAO1), strain MJ79 derived from PAO1-W (PMID 10984043), was provided by the Wargo Lab (Burlington, Vermont, USA). The MBEC Assay® 96-well plate was purchased from Innovotech Inc. (Edmonton, Alberta, Canada). Human dermal fibroblast cells, normal human adult (HDFa) were purchased from ATCC (Manassas, Virginia, USA). Fibrolast basal medium supplemented with mixture C-39315 was purchased from PromoCell (Heidelberg, Germany). The ReadyProbes™ Cell Viability Imaging Kit, Blue/Green was purchased from Thermofisher (Waltham, MA, USA). Ultrafiltered water was produced by Milli-Q Millipore Integral 10 water purification system. Filtered centrifuge tubes with a molecular weight cutoff of 100 kDa and regenerated cellulose membrane were purchased from Amicon (Merck KGaA, Darmstadt, Germany). Phosphate Buffered Saline (PBS) was purchased from Sigma Aldrich (St. Louis, Missouri, USA).

## Methods

### Synthesis of PDH-loaded alginate hydrogels

Alginate dressings are crosslinked ionically with divalent or polyvalent cations, such as calcium, magnesium, barium, zinc, or strontium, to form hydrogels. The use of calcium as the crosslinker adds hemostatic properties to the hydrogel [28]. In this study, Ca^2+^ ions crosslinked Na-Alg to entrap the enzyme PDH in hydrogels, as illustrated in Figure 1. Briefly, a buffer exchange process was used to remove glycerol from the supplied PDH and dissolve the enzyme in 50 mM MOPS at pH 7.4. The buffer-exchanged PDH solution was concentrated by centrifugation for 4 hours at 6500 rcf using filtered centrifuge tubes (molecular weight cutoff 100 kDa) and stored overnight in the refrigerator until use the next day. To prepare the hydrogels, varying concentrations and volumes of Na-Alg and CaCl_2_ (Table 1), based on similar hydrogels created for wound dressings [29-31], were prepared in ultrapure water. Based on the units of the enzyme to be entrapped in the hydrogel, a specific volume of the enzyme solution was added to the Na-Alg solution and gently mixed by pipetting up and down slowly to minimize bubble formation. The mixture was placed into molds and frozen at -20°C for 2 hours. Subsequently, CaCl_2_ was added dropwise to each mold. The mold was placed on a shaker (Fisher Scientific) with a speed of 20 tilts per minute for 30 minutes to facilitate crosslinking. After 30 minutes, to ensure an excess of Ca^2+^ was present for crosslinking throughout the thawing process, the solution was aspirated and replaced with fresh CaCl_2_ solution and then shaken for an additional 20 minutes. Finally, the remaining liquid was aspirated from the surface of the hydrogel, and the hydrogels were submerged in MOPS buffer to wash off any excess CaCl_2_. After washing with MOPS, the hydrogels were used for further studies.

**Figure 1.**
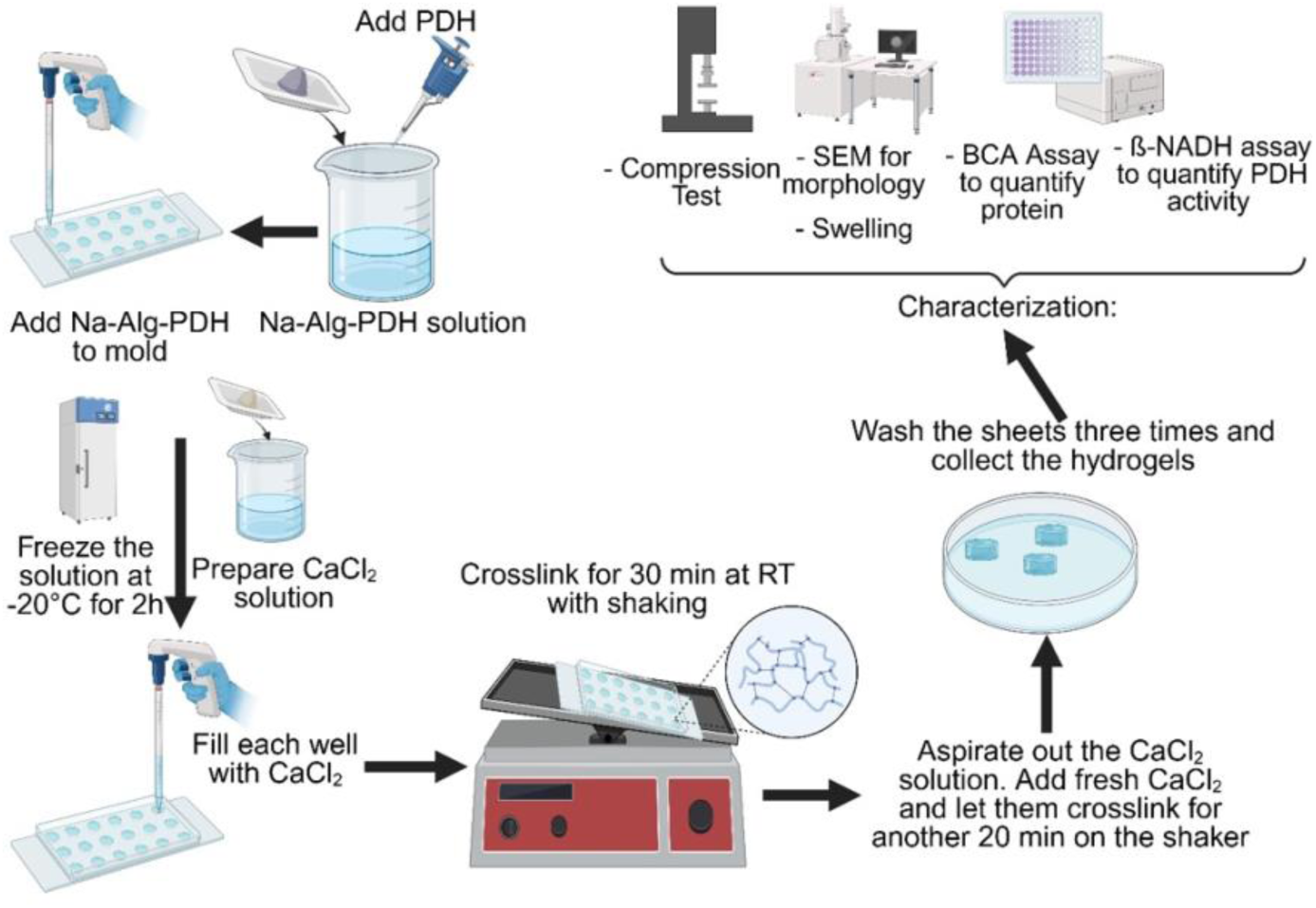
Synthesis and characterization of alginate-PDH Hydrogels. Created with BioRender.com

**Table 1.** List of alginate-PDH synthesis variables. All hydrogels were prepared with 2.5% Na-Alg and 0.4 M CaCl_2_.

| Na-Alg ($\mu\text{l}$) | PDH (U) |
| --- | --- |
| 720 | 7.198 |
| 720 | 10 |
| 240 | 2.9 |
| 240 | 3.4 |
| 65 | 0.05 |
| 65 | 0.1 |
| 65 | 0.2 |
| 65 | 0.35 |
| 65 | 0.475 |
| 65 | 0.6 |
| 65 | 0.725 |
| 65 | 0.85 |
| 65 | 0.975 |
| 65 | 1.1 |

### Unconfined Compression Testing

The compressive modulus of a hydrogel wound dressing is a crucial measure of its structural integrity and conformability to the wound, quantifying the amount of stress per strain in the elastic region of material deformation [32]. The alginate hydrogels without PDH were subjected to uniaxial compression immediately after synthesis while still hydrated. Upon formation in molds (0.59 × 0.59 inches), the hydrogels were punched into cylindrical shapes (8 mm diameter) to match the geometry of the compression platen. Unconfined compression tests were conducted at room temperature using a DHR-2 rheometer (TA Instruments, Delaware, USA) equipped with flat, stainless steel compression platens (8 mm diameter) and a Peltier plate. Hydrogel samples with heights less than half their diameter (h < 0.5Ø) were positioned on the Peltier plate, and the upper platen was lowered to apply a preload of 0.01–0.03 N. The initial gap was recorded as the original gauge length for strain calculation. Compression was applied at a deformation rate of 10 μm/s until the samples reached 20% compressive strain. GraphPad Prism was employed to create stress-strain curves from the force-displacement data. Compressive strength was determined as the stress corresponding to 15% strain, while compressive moduli were calculated from the linear region of the stress-strain curve. To calculate the compressive modulus, the slope of the linear region of the stress-strain curve was determined. Slopes were calculated for 10% strain intervals within the range of 2% to 20% strain [33, 34]. The first slope with an R² value greater than 0.98 (indicating a strong linear fit within that range) was selected to calculate the compressive modulus.

### Scanning Electron Microscopy (SEM)

Following crosslinking, the hydrogels were rapidly frozen using liquid nitrogen to preserve the internal structure. The frozen samples were lyophilized (Freezone 2.5, Labconco, MO, USA), cryo-fractured, and sputter-coated (Cressington 108, Ted Pella, Inc., CA, USA) with a 10 nm layer of Au–Pd for imaging. SEM (Zeiss Sigma 300 VP Field-Emission SEM, Oberkochen, Germany) was used to visualize the internal structure of the crosslinked hydrogel. All samples were prepared in triplicate, with 10 measurements taken from each replicate. This pore size data was analyzed using one-way ANOVA followed by Tukey’s post hoc test.

### Swelling and Weight Loss Study

Swelling and weight loss studies are crucial for assessing the fluid absorption capacity and structural stability of hydrogel wound dressings, which are essential for maintaining prolonged functionality [35]. In this hydrogel design, diffusion of pyruvate and cofactors into the hydrogel is required for reaction with the entrapped PDH enzyme; therefore, a high swelling ratio further enhances the performance of the dressing. Alginate hydrogels with a constant CaCl_2_ concentration of 0.4 M and varying Na-Alg (2%, 2.5%, and 3% (w/v)) were synthesized. The 0.4 M concentration of CaCl₂ was chosen because further increases do not affect the mechanical properties of alginate. Hydrogels were placed in a -80°C freezer overnight and subsequently lyophilized. The dry weights of the hydrogels were measured and recorded as W_initial dry._ Next, each hydrogel was submerged in 4 ml of simulated wound exudate fluid for various periods of time (0.5, 1, 2, 3, 4, 8, 12, 24, 36, and 54 hours), and the swollen weights were recorded as W_swollen_. The simulated wound exudate fluid contained 0.02 M calcium chloride, 0.4 M sodium chloride, 0.08 M tris-methylamine, and 2% (w/v) bovine serum albumin (BSA) dissolved in ultrapure water [35]. The swelling ratio and water content were calculated using Equations 1 and 2, respectively [36, 37].

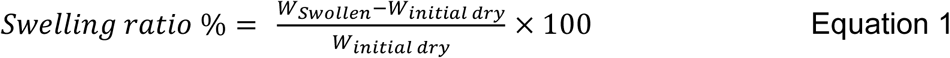

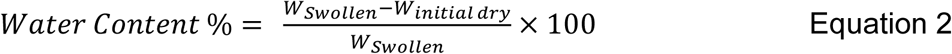

After measuring the swollen weights, the samples were dried again, and their final dry weights were recorded. The weight loss percentage was then calculated using Equation 3 [38, 39].

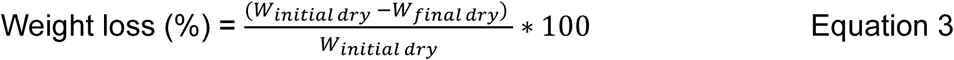

### Water Retention Capacity

Wound dressings need to retain absorbed liquids even when subjected to external forces. One way to apply uniform force to the hydrogels and assess their ability to retain liquid is by using centrifugation [40, 41]. In this method, all hydrogels were initially dried according to the protocol described in the swelling study and then allowed to swell in water at room temperature for 24 hours. The dry weight was recorded as W_d_. The weight of the swollen samples was measured (denoted as W_s_), and then they were centrifuged at various speeds (2000, 8000, and 16000 g) at room temperature. The weight after centrifugation was recorded as W_c_. The water retention capacity was then calculated using the following equation:

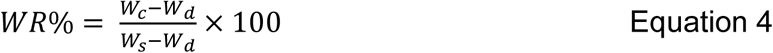

### Gelation Kinetics

Rheometry was conducted using an AR2000 rheometer (TA Instruments, New Castle, DE, USA) with a Peltier plate to determine the viscosity of alginate solutions before gelation as well as the kinetics of gelation in ionically crosslinked alginate. These analyses were carried out at 37°C, using a 20-mm diameter 1°59’6” steel cone geometry with a 57 µm gap between the plate and the cone. The polymer solutions were placed on the rheometer plate, and the cone was adjusted until an axial force was detected. A range of shear rates, from 1 to 100 (1/s), was applied for 120 seconds to measure the viscosity (n = 4), which was then computed and graphed using TA Instruments Data Analysis software. To study the gelation kinetics of these precursor solutions, measurements were performed at a frequency of 1 Hz and with a 1% oscillatory radial strain. For all samples, a 0.4 M solution of CaCl_2_ was sprayed on the sample half a minute into the test.

### Biochemical Assays: Enzyme Activity and Protein Quantification

PDH activity was measured using an assay to quantify the conversion of β-NAD^+^ to β-NADH, detected spectrophotometrically at 340nm over various time periods. This assay, referred to as the PDH activity assay, enabled the calculation of PDH activity (mU/ml) for each sample using an extinction coefficient of 6.22 mol^-1^cm^-1^ at 340 nm. Assays were conducted at room temperature in a total volume of 1 ml, including the following cofactors: 2 mM β-NAD^+^, 2 mM CoA, 20 μM TPP, 50 μM MgSO_4_, and 10 mM sodium pyruvate dissolved in MOPS. At each time interval, 200 µl of the solution was aliquoted into a 96-well plate, and the absorbance was measured at 340 nm using a SpectraMax™ i3x microplate reader (Molecular Devices, USA).

The amount of protein entrapped within each hydrogel was measured using a Micro BCA™ Protein Assay Kit. First, the hydrogels were dissolved in a 16.7 mM sodium citrate solution under gentle rotation (Tube Revolver, Thermo Scientific, MA, USA) for 48 to 72 hours, depending on the size of the hydrogels. Subsequently, the assay was conducted following the protocol provided by the manufacturer. Briefly, 150 μl of each sample was mixed with 150 μl of BCA working reagent, incubated for 2 hours at 37°C, and the absorbance was read at 562 nm. The concentration was then calculated using the albumin standard curve, prepared with water as the solvent.

### Retention of Enzyme Activity After Storage

To evaluate the storage stability of the PDH when entrapped in alginate hydrogels, three aliquots of unencapsulated PDH and three samples of PDH-loaded hydrogels were submerged in a large volume of 50 mM MOPS buffer (pH 7.4) and incubated at 4 °C. Samples were collected after 24, 48, 96, and 144 hours of incubation. All samples were continuously agitated, and aliquots were taken to perform both the PDH activity assay and the BCA protein assay.

### PDH Release Kinetics

Each hydrogel sample was placed into a separate 15 ml conical, and 10 ml of MOPS buffer was added to initiate the release study. This experiment was performed at 35°C to simulate a topical wound environment [42]. To maintain sink conditions, the total volume of the release medium was significantly higher than the sample volume. The conicals were then placed on a shaker and agitated gently. For the first hour, samples were collected every 20 minutes, followed by every 30 minutes for the next 5 hours, and then at 8, 10, and 12 hours. At each time point, two aliquots of 20 µl each were collected for the PDH activity assay and two aliquots of 150 µl each for the BCA assay. To measure the activity of the hydrogels after the release study, each sample was removed from the MOPS solution and submerged in 1 ml of PDH working solution. Each sample was placed on a tube rotator for 60 minutes. Then, 200 µl of each sample was transferred to a 96-well plate, and the absorbance was measured at 340 nm.

### Biofilm Culture

*Pseudomonas aeruginosa* (PAO1), generously donated from the Wargo lab, was stored at -80°C in media containing 10% skim milk. Bacteria were scraped and incubated in LB broth overnight at 37°C. The OD_600_ of the overnight culture was adjusted to 0.1 with 0.85% saline. Then, 150 µl of the 10-fold diluted LB broth was added to each well of a minimum biofilm eradication concentration MBEC Assay® 96-well plate and incubated overnight at 37°C and 110 rpm to form biofilms on the pegs. Media contained 2 mM β-NAD^+^, 2 mM CoA, 20 µM TPP, and 50 µM MgCl_2_ in 10-fold diluted LB. Negative control samples of bacteria alone lacked cofactors. PDH-loaded hydrogels and 150 µl of media were added to each well. Alginate hydrogels containing heat-inactivated PDH (HI-PDH) were also tested as a control. The biofilms grown on the pegs were then placed into the wells containing the media with hydrogel samples at the bottom of the well and incubated for 24 hours at 37°C and 110 rpm. After incubation, the biofilms were stained with 1% CV dye, air-dried, placed in 150 µl of 95% ethanol, and the absorbance of the resulting CV solution was measured at 570 nm.

### Cell Culture

Primary normal human adult dermal fibroblasts (HDFa) were seeded onto 96 well plates at a density of 7,200 cells per cm^2^ and grown in fibroblast growth media, supplemented with 1 ng/ml basic fibroblast growth factor (recombinant human) and 5 µg/ml insulin (recombinant human) at 37°C with 5% CO_2_. To quantify the cell viability in the presence of hydrogels, the Blue/Green ReadyProbes^TM^ kit was used to determine the percentage of living cells. Following ISO 10993-5:2009, hydrogels (n=3) were washed twice with 3 ml of PBS and twice with 3 ml of cell media before suspension in cell media for 24 hours to obtain hydrogel leachate [43-46]. After 24 hours, the leachate was diluted 1:2 nine times in cell media and added to confluent cells. Controls were 4% Triton X-100 and untreated cells exposed to media alone. After 24 hours of incubation with cells, the leachate was aspirated, and NucBlue and NucGreen dyes, combined with cell media, were added to each well for 30 minutes before measuring at the respective fluorescence excitation and emission wavelengths

### Statistical analysis

GraphPad Prism version 10.2.2 (GraphPad Software, CA, USA) was utilized for all statistical analyses. Unless otherwise noted, data were collected in triplicate and analyzed using two-way ANOVA followed by Tukey’s multiple comparison test, with statistical significance set at p < 0.05. The biofilm and cell viability assays were analyzed using a one-way ANOVA followed by Tukey’s post hoc t-test, with statistical significance set at p < 0.05, was performed with n=3 sample size in GraphPad Prism.

## Results and Discussion

### Optimizing Hydrogel Stiffness: Impact of Alginate and Calcium Concentrations

The compressive modulus of a wound dressing is linked to patient comfort and the support of healing [47]. The compressive modulus of a hydrogel wound dressing is a vital indicator of its structural integrity and ability to conform to a wound, representing the stress-to-strain ratio within the material’s elastic deformation range [32]. Unloaded alginate hydrogels were tested under uniaxial compression immediately after synthesis, while still in their hydrated state in order to determine which formulation generated the most comparable stiffness to commercially available hydrogels. Alginate hydrogels crosslinked with Ca^2+^ ions exhibit a compressive modulus ranging from 1 to 100 kPa, depending on the concentrations of alginate and crosslinker [48, 49]. Hydrogels used in wound dressing applications typically have compressive moduli ranging from 15 to 90 kPa [50-53]. Figure 2 shows the effects of Na-Alg and CaCl_2_ concentrations in A and B, respectively. As shown in Figure 2B, increasing the Na-Alg concentration from 2% to 3% results in a significant rise in compressive modulus across all samples. This trend is attributed to the formation of a denser and more interconnected polymer network, which enhances the stiffness and resistance to deformation of the hydrogel [54, 55]. The modulus of the 2.5% sample falls between the values for 2% and 3%. Figure 2A shows that increasing the CaCl₂ concentration from 0.2 M to 1 M does not substantially impact the compressive modulus except in hydrogels with the highest Na-Alg concentration (3%), suggesting that at lower Na-Alg concentrations, 0.2 M CaCl₂ is sufficient for effective crosslinking. However, at 3% Na-Alg, the addition of more CaCl₂ enables further crosslinking, resulting in a more robust and tightly crosslinked network with a higher compressive modulus. Notably, 0.4 M CaCl₂ is the threshold concentration at which further increases do not significantly enhance the compressive modulus in any of the tested samples. Therefore, 0.4 M CaCl₂ was selected for use in subsequent studies.

**Figure 2.**
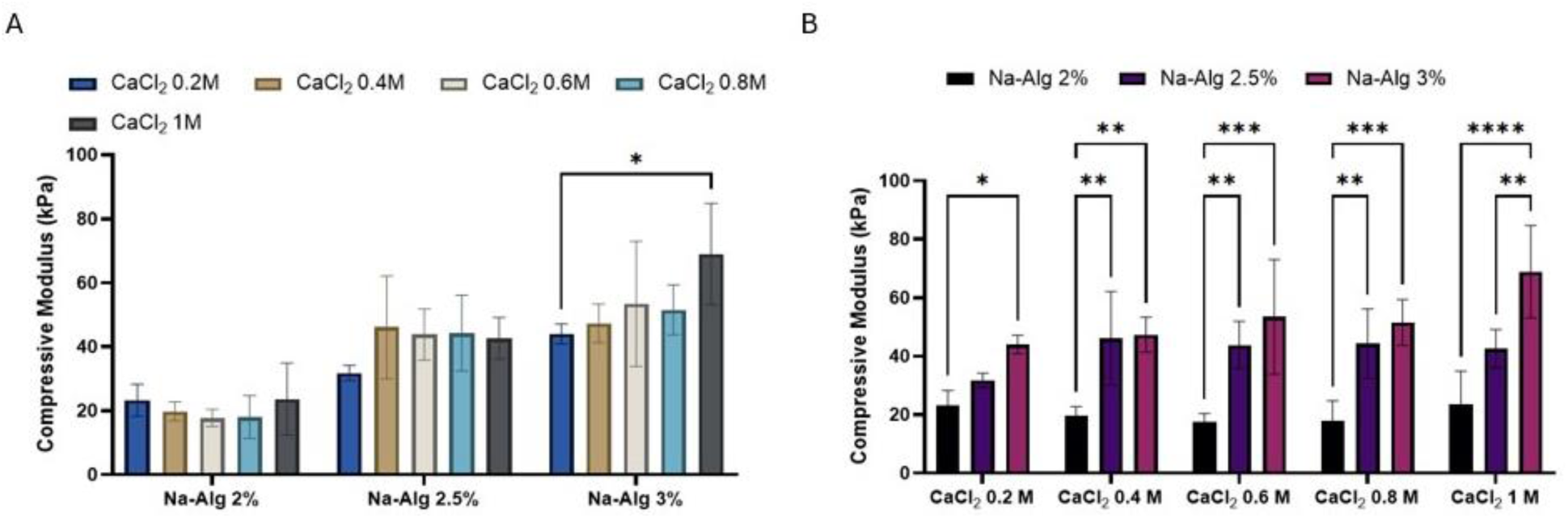
Compressive modulus of varying concentrations of Na-Alg and calcium chloride. A) Impact of calcium chloride concentration (0.2-1 M) on compression modulus of synthesized hydrogels, B) Impact of Na-Alg concentrations (2, 2.5, and 3%) on compression modulus of synthesized hydrogels. Data shown are mean ± standard error of the mean. n=3, *p<0.05, **p<0.005, ***p< 0.0003, ****p< 0.0001.

### Porous Microstructure of Alginate Hydrogels Enables Diffusion of Small Molecules for Enzyme Interaction

SEM imaging was employed to examine the internal structure of the unloaded alginate hydrogels and assess the impact of Na-Alg concentration on pore morphology. The hydrogel was designed to allow diffusion of small molecules such as pyruvate into the hydrogel matrix, where they interact with the encapsulated PDH, followed by diffusion of the products out of the matrix. Therefore, a porous structure with interconnected pores was essential to facilitate efficient small molecule transport while restricting movement of the large enzyme. Using SEM, images of the internal and external structures of the hydrogels were captured for the lowest (2%) and highest (3%) concentrations of Na-Alg, as shown in Figure 3E. The observed pore size range was consistent with previously reported data on alginate hydrogels at either Na-Alg concentration [33, 56]. PDH is a relatively large enzyme, with a diameter of approximately 45 nm [57]. A porous structure was essential for high encapsulation efficiency and to enable cofactors and pyruvate to diffuse while the enzyme was entrapped. The SEM images indicated that a porous, interconnected network was created within the alginate hydrogels, with pore sizes suitable for allowing the movement of small molecules through the matrix.

**Figure 3.**
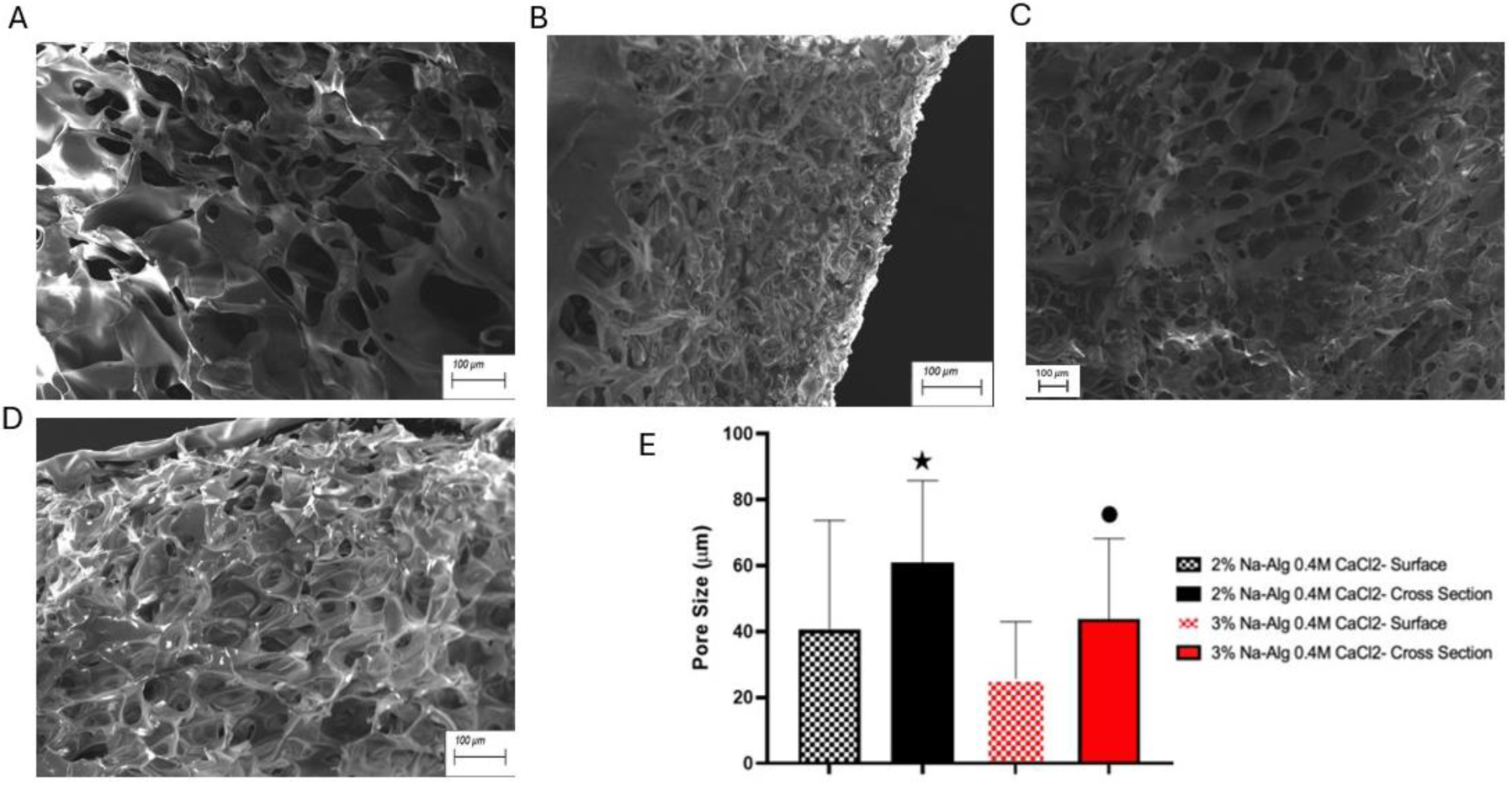
SEM images of the following samples: A) the external surface of 2% Alg-0.4M CaCl_2_ hydrogel, B) the external surface of 3% Alg-0.4M CaCl_2_ hydrogel, C) the cross-sectional surface of 2% Alg-0.4M CaCl_2_ hydrogel, and D) the cross-sectional surface of 3% Alg-0.4M CaCl_2_ hydrogel. E) Analysis of impact of varying Na-Alg concentration on hydrogels’ pore size by cross-sectional and surface imaging. Data shown are mean ± standard error of the mean. Number of hydrogel samples made from each composition = 3, number of measured pores from each sample 10, total of 30 measurements from each composition. ⬤p<0.01, *p< 0.0001 compared to 3% Na-Alg 0.4 M CaCl_2_ – Surface.

### Swelling and Water Retention in Hydrogels

A high swelling ratio is particularly crucial in this hydrogel system to enhance the diffusion of pyruvate and cofactors into the system, allowing them to react with the entrapped PDH. Furthermore, the swelling ratio is a measure of wound dressing efficacy in managing exudate absorbance, which promotes faster healing and reduces the risk of maceration. Figure 4A illustrates the swelling ratios of alginate hydrogels with varying concentrations of Na-Alg (2%, 2.5%, and 3%). The swelling behavior in the presence of simulated wound exudate fluid was time-dependent, with a rapid increase during the first 4 hours, followed by a slower absorption rate that stabilized after 12 hours. There was a statistically significant decrease in swelling behavior as the concentration of Na-Alg increased from 2% to 3%. A similar trend was observed in the water content percentage (Figure 4B). Both swelling behavior and water content percentage inversely correlate with crosslinking density, as hydrogels with tighter networks exhibit lower swelling ratios and reduced water content [37]. Following this trend, samples with higher Na-Alg concentrations demonstrated a greater ability to absorb water and retain it effectively compared to those with lower concentrations. The water content capacity of all samples ranged from 74% to 94%, meeting the standard requirements for wound dressings [36].

**Figure 4.**
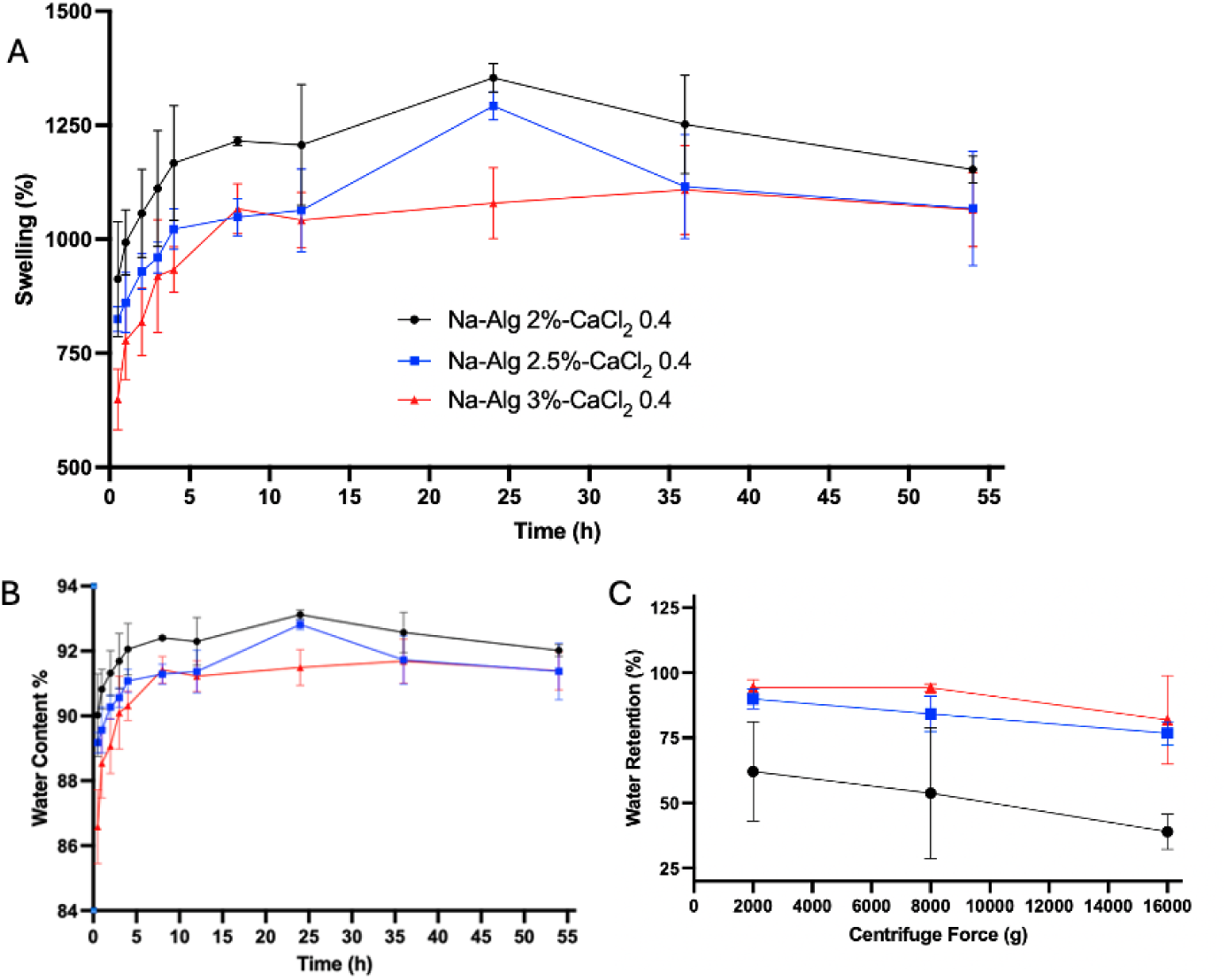
A) Swelling ratio percentage of hydrogels in water calculated by equation 1, B) water content percentage calculated by equation 2, and C) water retention percentage of varying concentrations (2, 2.5, and 3%) of Na-Alg crosslinked with 0.4 M CaCl_2_ over 54 hours compared to the initial dry weight. Data shown are mean ± standard error of the mean. Hydrogels were prepared in triplicate (n=3).

The water retention capacity in the presence of simulated wound exudate fluid, shown in Figure 4C, further supports these findings. Among the tested concentrations, the sample with 2% Na-Alg exhibited a significantly lower capacity to retain water within its structure.

There was no statistically significant change in water retention percentage when the Na-Alg concentration increased from 2.5 to 3%. This effect can be attributed to the hydrogel’s lower crosslinking density and larger pore sizes, which allowed water to escape more readily. These results highlight the importance of optimizing Na-Alg concentration to achieve desirable swelling and retention properties for successful wound dressing applications.

### Structural Stability of Alg Hydrogels in Water and Simulated Wound Fluid Vary Slightly with Alginate and Calcium Concentration

In this hydrogel, minimal weight loss was essential to maintain the structural integrity of the PDH-loaded hydrogel, ensuring its stability and functional performance in the presence of water and wound exudate. After an initial weight loss within the first 24 hours, no further significant changes were observed over the subsequent three days for any sample, except for the 2% Na-Alg hydrogel (Figure 5). This stability in the hydrogel structure is advantageous for this wound dressing application, as it ensures the integrity required to retain the enzyme within the hydrogel matrix while allowing the diffusion of pyruvate and wound exudate into the hydrogel. In clinical settings, alginate hydrogels are used to treat wounds that produce moderate to heavy exudate while the dressing is changed every one to three days [28]. These data indicated that the 2.5% and 3% samples did not exhibit significant structural degradation over three days, making them suitable for use in medical applications.

**Figure 5.**
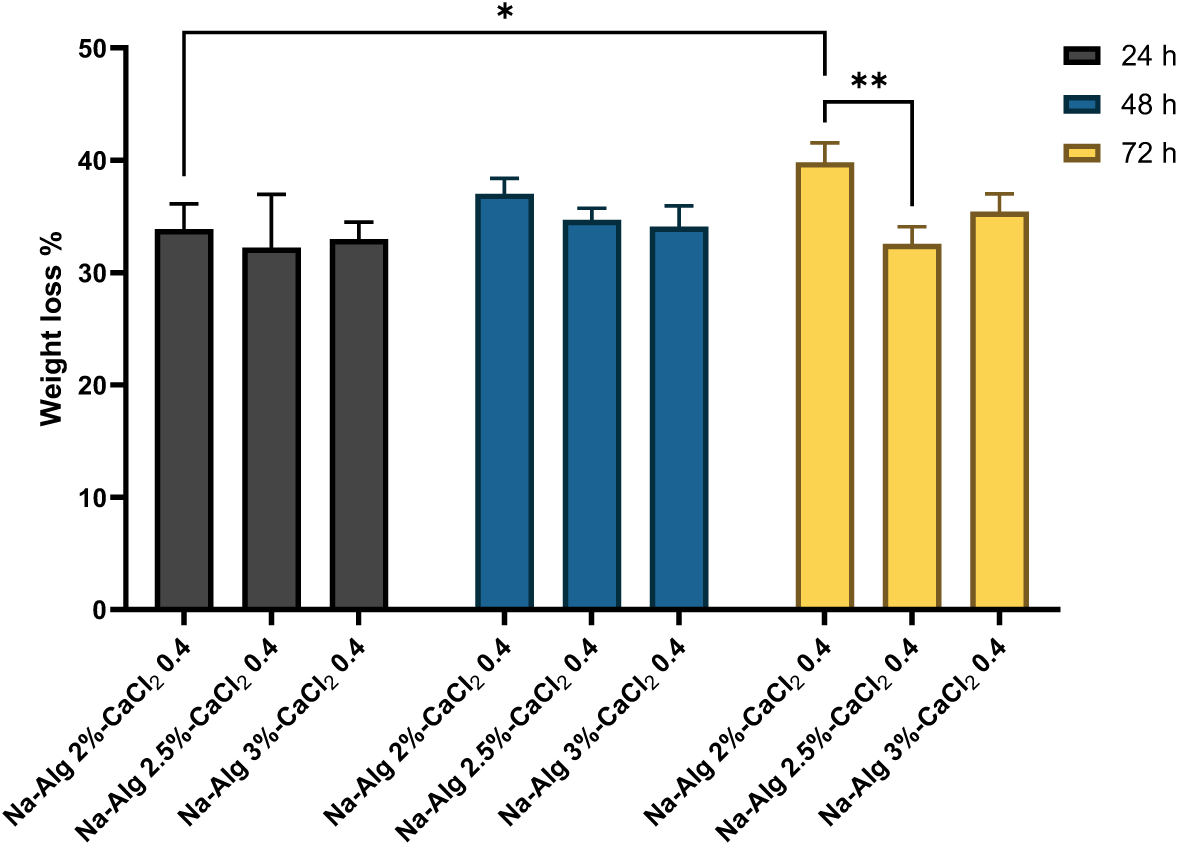
Weight loss study performed in ultrapure water on 2, 2.5, and 3 % Na-Alg crosslinked with 0.4 M CaCl_2_ hydrogels after 24, 48, and 72 hours. Data shown are mean ± standard error of the mean. Hydrogels were prepared in triplicate (n=3). *p < 0.04 and **p <0.003.

The high salt concentration and moist environment of wounds can disrupt the hydrogel’s polymer network, leading to structural degradation. To assess the swelling behavior of the hydrogels in a simulated exudate containing salt and protein, a more physiologically relevant solution than water, hydrogels were exposed to wound simulated fluid (WSF).

When exposed to WSF, as shown in Figure 6, all hydrogels experienced negative weight loss. This outcome was attributed to the absorption of salts by the hydrogels, which increased their weight, resulting in a negative weight loss. Notably, the hydrogel with 2% Na-Alg exhibited a significant weight decrease from 24 to 48 hours, suggesting reduced structural stability in the simulated wound exudate. In contrast, hydrogels with higher Na-Alg concentrations maintained their weight over the same period, indicating better structural integrity.

**Figure 6.**
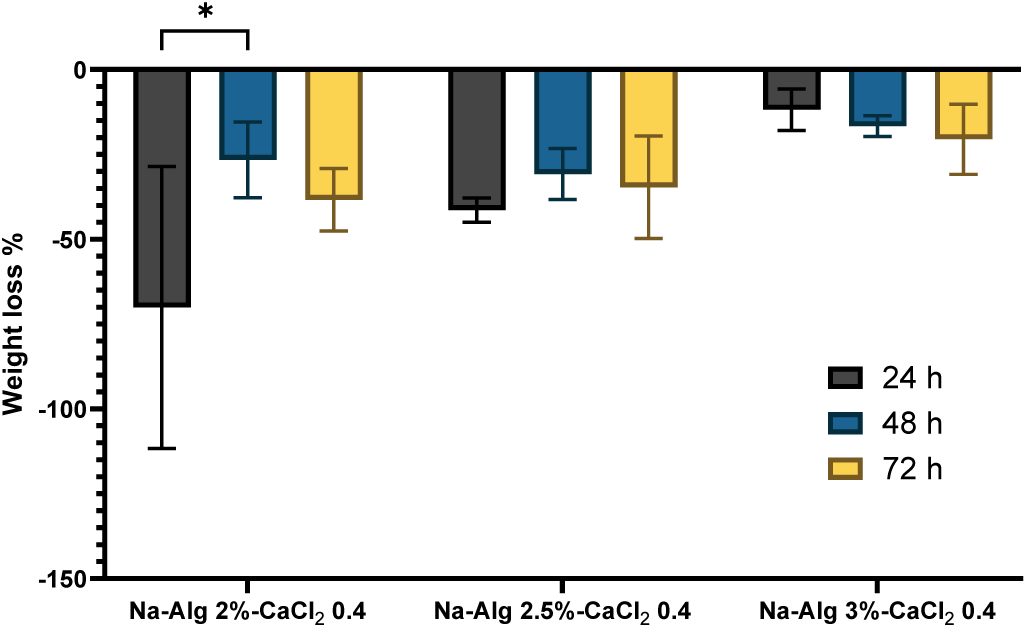
Weight loss in WSF on 2, 2.5, and 3 % Alg hydrogels crosslinked with 0.4 M CaCl_2_ after 24, 48, and 72 hours. Data shown are mean ± standard error of the mean. Hydrogels were prepared in triplicate (n=3), *p < 0.02.

### Alginate Concentration-Dependent Viscosity and Rapid Gelation Kinetics

Viscosity and gelation are critical for hydrogels as they determine their ease of application and directly impact the hydrogel’s performance in drug delivery and wound healing by ensuring structural integrity and functionality. Figure 7 shows the viscosity versus shear rate curves for 2%, 2.5%, and 3% Na-Alg solutions. Viscosities remained relatively consistent despite increasing shear rates, highlighting the non-Newtonian nature of Na-Alg solutions. The viscosity exhibited a concentration-dependent increase. For instance, at a shear rate of 57.33 s⁻¹, the viscosity values for 2%, 2.5%, and 3% Na-Alg were 0.0492, 0.0547, and 0.0811 Pa·s, respectively. This trend aligns with previous studies that report enhanced viscosity with higher polymer concentrations [58]. The findings align with the polymer solution theory, which suggests that higher polymer concentrations lead to increased intermolecular interactions, thereby enhancing viscosity [59].

**Figure 7.**
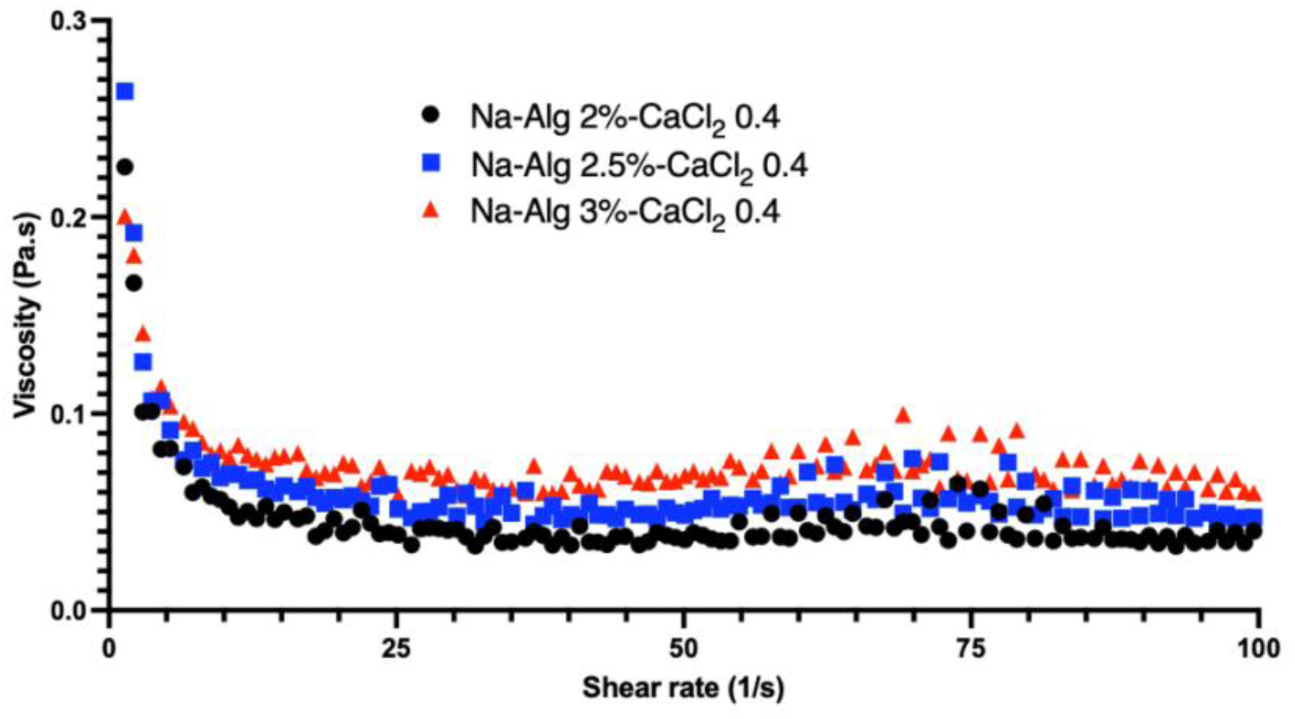
Viscosity values versus shear rate at 2, 2.5, and 3% Na-Alg concentration of hydrogels. (n=1.)

Figure 8 illustrates the gelation kinetics of Na-Alg solutions. Initially, within the first 30 seconds and before the addition of the crosslinker CaCl_2_, the loss modulus (G") exceeds the storage modulus (G’), indicating a predominantly liquid phase with viscous behavior. Upon the introduction of the crosslinker, there is a substantial increase in both G’ and G", suggesting the onset of gelation. According to the literature, the gelation time of ionically crosslinked alginate is typically on the order of a few seconds [60]. In this study, all samples exhibit short gelation times (31–33 seconds), consistent with the rapid gelation behavior characteristic of alginate systems.

**Figure 8.**
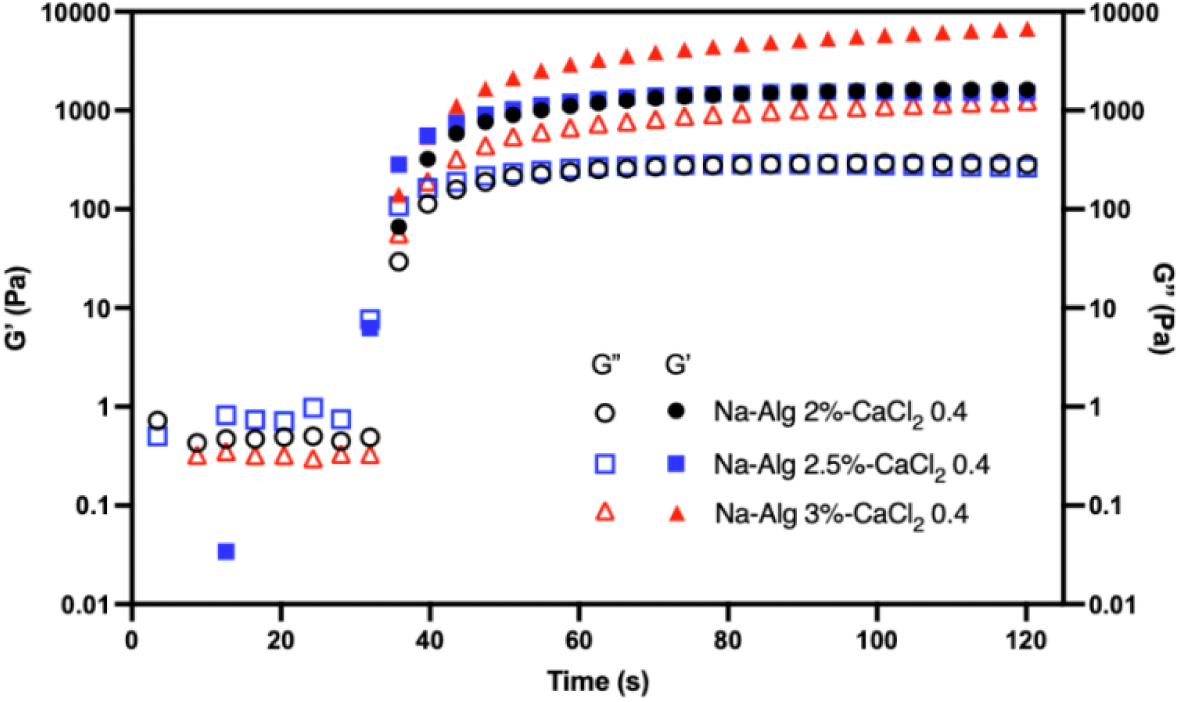
Representative curves of storage (G’) and loss (G’’) moduli for ionically crosslinked alginate hydrogel. (n=1).

Based on the gathered data, 2.5% Na-Alg and 0.4 M CaCl_2_ were selected for further analysis. These results for compressive moduli, weight loss percentage, and water retention indicated that 2% Na-Alg hydrogels exhibited significantly weaker mechanical and physical properties compared to higher concentrations. Additionally, 3% Na-Alg solutions were highly viscous (Figure 8), which was compounded by the already viscous enzyme solution. To dissolve the components, vigorous mixing by vortex was required, which resulted in a loss of enzymatic activity [61]. Additionally, the high viscosity of the 3% Na-Alg solution made it difficult to mold the samples effectively. Therefore, 2.5% Na-Alg was determined to be the optimal choice for further development of PDH-encapsulating hydrogels.

### PDH-Loaded Hydrogels Achieve Threshold Activity for Biofilm Dispersion in Hydrogel Size-Constrained Matter

Since the depletion of pyruvate disrupts bacterial metabolism and promotes biofilm dispersion, assessing the enzyme’s activity when entrapped in the hydrogels was critical. Previously published work from our group shows that the presence of 5 mU of unencapsulated enzyme resulted in a 2.2-fold reduction in biofilm biomass, while the addition of 10 and 20 mU of PDH resulted in a 2.9-fold reduction [26]. *In vitro* and *in vivo* studies indicate that 5 mU of unencapsulated PDH has sufficient enzymatic activity to disperse biofilms [62]. In the present study, 5 mU of unencapsulated PDH was used as a baseline comparison to the encapsulated samples. Figure 9 displays the activity of various PDH-encapsulating hydrogels compared to unencapsulated PDH over 60 minutes. The initial studies were conducted on hydrogels formed in round molds with a diameter of 8 mm and a volume of 240 μl, as well as hydrogels molded with a surface area of 15x15 mm with a total volume of 720 μl. These data show that samples 720 μl Alg + 10 U PDH, 240 μl Alg + 2.9 U PDH, and 240 μl Alg + 3.4 U PDH show the same level of activity as the 5 mU unencapsulated PDH, meeting the threshold for antibiofilm activity [26].

**Figure 9.**
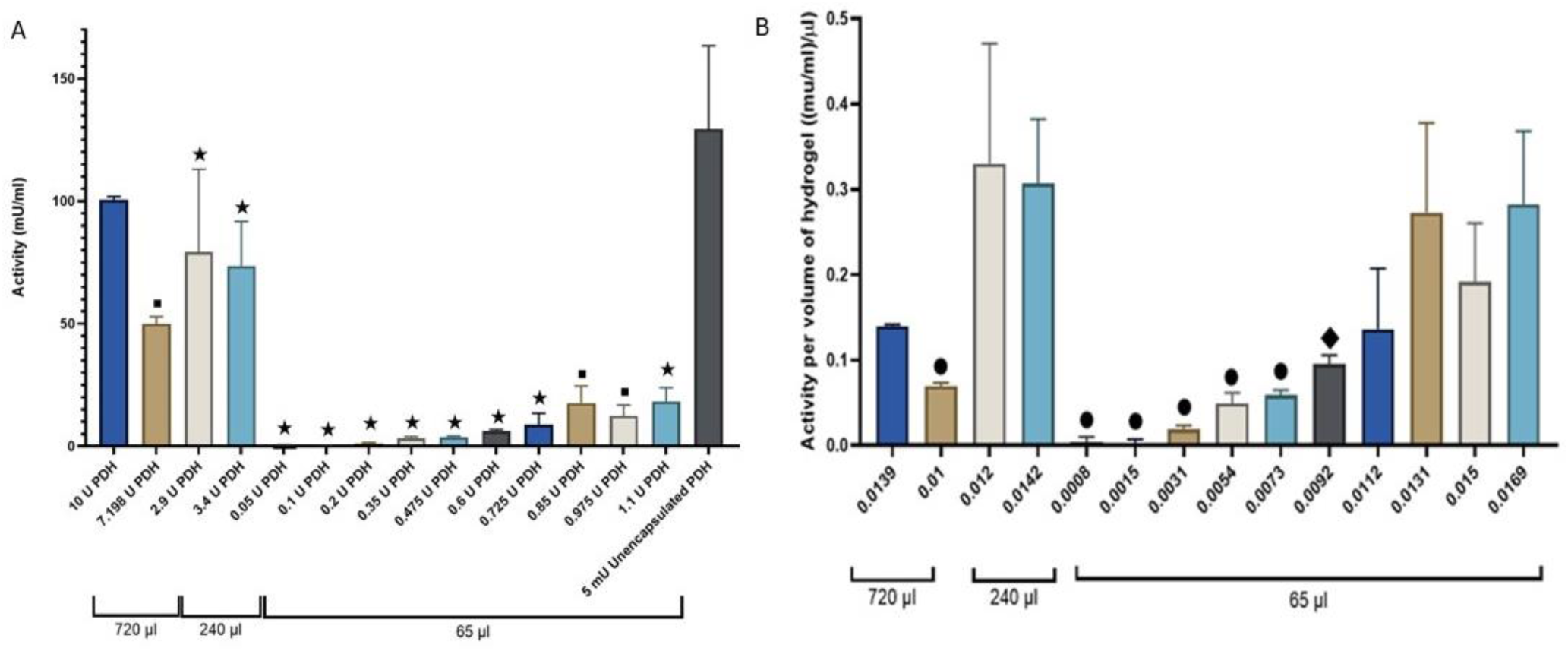
A) PDH activity of PDH-loaded alginate hydrogels with sizes of 720 µl, 240µl and 65µl and values of 0.05, 0.1, 0.2, 0.35, 0.475, 0.6, 0.725, 0.85, 0.975, 1.1, 2.9, 7.198, and 10 U loaded PDH. B) PDH activity normalized per the volume of hydrogels vs. the initial U PDH of loaded enzyme per µl of hydrogel (theoretical loading). Data shown are mean ± standard error of the mean. Hydrogels were prepared in triplicate (n=3). Statistically significant comparisons denoted as ▪p<0.001, *p< 0.0001 compared to unencapsulated PDH. Statistically significant comparisons denoted as ⬤p<0.001, ♦p<0.01 compared to Alg 65 µl + 0.0131 U PDH.

To conduct biofilm assays on the MBEC Assay® platform, smaller hydrogels of 65 μl volume were necessary. The activity study demonstrates that when the alginate hydrogel volume is 65 μl, the enzymatic activity is significantly lower than that of unencapsulated PDH. At concentrations below 0.35 U, the PDH encapsulated within the hydrogels shows minimal activity. However, the activity increases with enzyme concentration, reaching a plateau beyond 0.85 U of encapsulated PDH. This concentration-dependent trend contrasts with previous studies encapsulating PDH in PLGA nanoparticles, where increasing the loaded PDH led to fluctuations in activity, and a sharp drop was observed after reaching peak activity [27]. The variability in the activity of the NPs suggests that alginate hydrogels offer a more predictable and reliable material for future use. Figure 9B displays the activity of PDH normalized by the volume of the hydrogel. Although direct comparison of the different size hydrogels with matched theoretical loading was not possible due to variations in the concentration of the enzyme, enzyme activity appears to depend somewhat on hydrogel size. For example, when theoretical loading is 0.013 U/μl, activity differs between small and large hydrogels, implying that diffusion-based transport limitations should be investigated further.

In previous studies when PDH was encapsulated in PLGA NPs, activity increased from 1.9 to 30.2 mU/ml as the loaded PDH increased from 3.488 U to 6.96 U [27]. In the current hydrogel system, the highest activity observed was 17.7 ± 6.86 mU/ml at 0.85 U of loaded PDH. Although this value is lower than expected, due to the size constraints of the well plate preventing an increase in hydrogel volume, this formulation was selected for further study. In other studies, alginate hydrogels loaded with enzymes such as β-galactosidase and maltogenic amylase have demonstrated higher enzymatic activities of 0.65 IU/mL and 17.0 U/mg, respectively. However, it is important to note that these enzymes and their catalytic reactions differ significantly from PDH. More importantly, those systems were designed for enzyme release, allowing the enzymes to diffuse and freely interact with cofactors in the surrounding medium, thereby maximizing catalytic efficiency. In contrast, in our system, PDH remains entrapped within the hydrogel matrix, and the reaction depends on the passive diffusion of cofactors and pyruvate into the hydrogel. The enzymatic conversion occurs within the confined environment of the hydrogel, and substrate exchange is limited to the rate of diffusion and replenishment from the surrounding solution [63, 64].

### Effect of PDH Loading on Encapsulation Efficiency is Insignificant Above 0.2 U

Encapsulation efficiency is essential for hydrogels as it determines how effectively bioactive agents, like PDH, are incorporated and retained within the gel. The encapsulation efficiency (EE%) was measured for various concentrations of entrapped PDH inside the alginate hydrogels. Figure 10 shows an inverse relationship between the amount of PDH added and the EE% within the alginate matrix. At lower enzyme concentrations, particularly 0.05 U and 0.1 U PDH, the EE% is significantly higher, reaching approximately 35%. This elevated efficiency at low PDH loadings is attributed to the availability of free volume within the alginate network, which facilitates the effective entrapment of the enzyme. However, as the PDH concentration increases beyond 0.1 U, a decline in EE% is observed, likely due to limited free volume within pores. Moreover, at higher enzyme concentrations, PDH may exhibit aggregation behavior or interfere with the gelation dynamics of the alginate matrix, reducing its ability to become efficiently entrapped. Compared to other studies, the EE% for PDH in alginate is significantly lower than enzymes such as β-galactosidase and acetylcholinesterase, which have been shown to be encapsulated at 72% and 68%, respectively [64, 65]. Both β-Galactosidase and acetylcholinesterase are twenty times smaller than PDH, so their high EE% is likely due to this size disparity. Despite this decrease in EE%, activity assays reveal that functional enzyme activity continues to increase with higher PDH loading, reaching a plateau at 0.85 U. Therefore, from both an efficiency and activity standpoint, 0.85 U may represent the most practical upper limit for enzyme loading in this alginate-based delivery system. In the study on the encapsulation of PDH into PLGA NPs, the EE% was 11.5 ± 6.0%, with no established trend observed between the concentration of PDH and the EE%. These results suggest that the hydrogel may serve as a more reliable material for encapsulating PDH compared to PLGA NPs.

**Figure 10.**
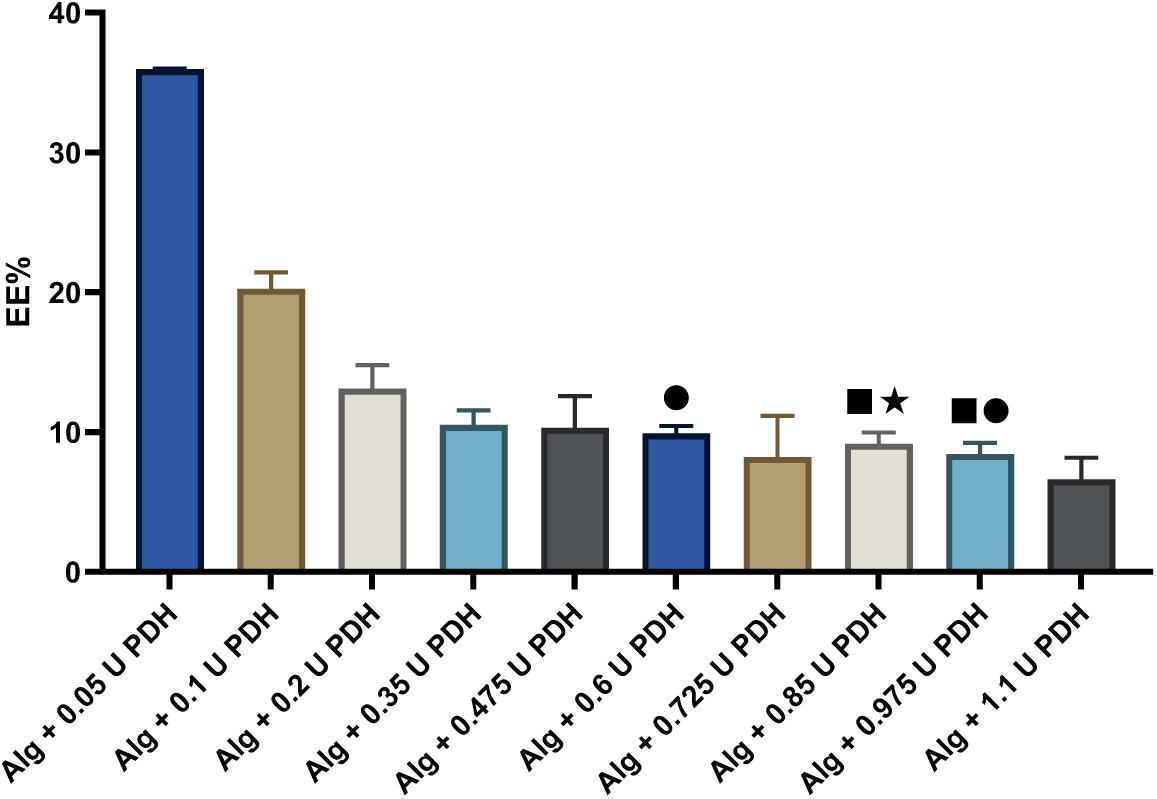
EE% of PDH in amounts of 0.05, 0.1, 0.2, 0.35, 0.475, 0.6, 0.725, 0.85, 0.975, and 1.1 U in 65μl alginate hydrogels. Data shown are mean ± standard error of the mean. Hydrogels prepared in triplicate (n=3). ⬤p<0.01 and *p<0.001 compared to Alg hydrogels containing 0.05 U PDH. ▪p<0.01 compared to Alg hydrogels containing 0.1 U PDH.

### Alginate Encapsulation Preserves PDH Activity with Gradual Decline Over Time

Retaining enzyme activity over time is crucial to ensure sustained antibacterial properties. The activity of alginate-entrapped PDH was compared to that of unencapsulated PDH stored at 4 °C under matched conditions to evaluate the activity after storage over six days. Figure 11 displays the activity of both samples, which declines over time. The loss of activity in the entrapped sample was more pronounced than in the unencapsulated PDH. However, previous work with PLGA-entrapped PDH, demonstrates a rapid initial decrease in activity with activity falling to below 30% within the first 50 hours [27]. In contrast, the alginate-based system retained almost full activity during the first 24 hours, with a gradual decline thereafter, retaining approximately 60% of its initial activity after six days. Interestingly, while the PLGA system showed some recovery in activity over time, the alginate formulation demonstrated a more predictable and consistent loss of activity.

**Figure 11.**
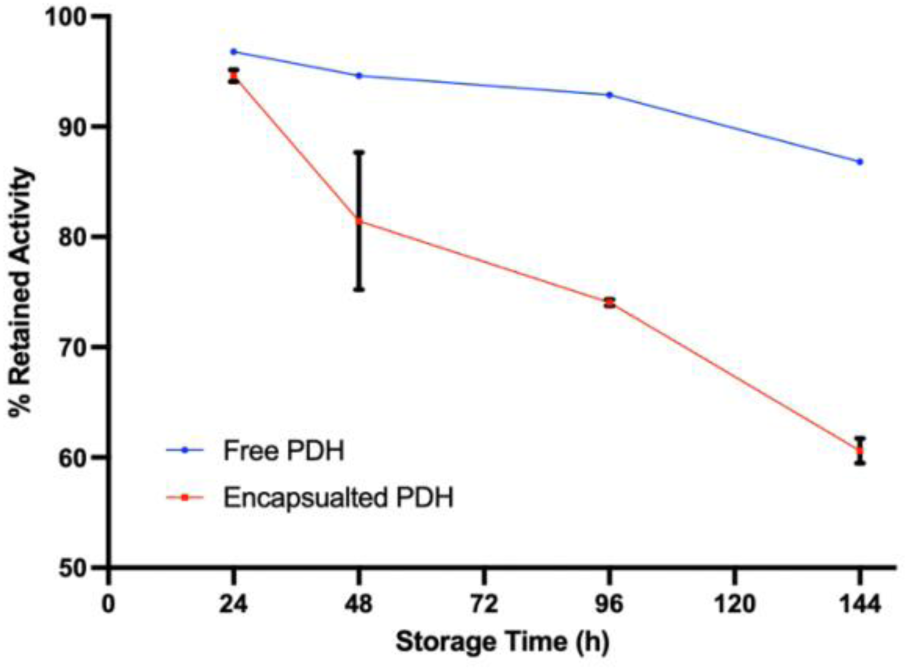
Percentage of retained activity of PDH loaded in alginate hydrogels compared to unencapsulated PDH after incubation at 4 °C. Data shown are mean ± standard error of the mean. Hydrogels prepared in triplicate (n=3).

### Most PDH Is Retained Alginate Hydrogels for >12 hours

Retaining enzymes within the hydrogel is vital to ensure sustained activity and localized functionality over time. Cumulative enzyme release and corresponding activity are presented in **Error! Reference source not found.**. Importantly, as shown in Figure 12A, after 12 hours of incubation in buffer, 28.01 ± 13.43% of the entrapped enzyme was released from the hydrogel, indicating that most of the enzyme is retained over a time period suitable for the wound dressing to be used in practice, as desired. Figure 12B demonstrates that the released fraction of the enzyme exhibited no enzymatic activity over the entire release period. In contrast, the activity of the hydrogels themselves after 12 hours of incubation in the stated conditions was still 48.97 ± 4.62% of the initial activity. The percentage of enzymatic activity relative to the initial activity showed that the activity retained within the hydrogels after incubation was significantly higher than that of the released enzyme at each time point during the 12-hour period (p < 0.0001).Together, these results validate the intended design of this delivery system whereby PDH is entrapped within the hydrogel while pyruvate and cofactors diffuse into the polymeric matrix, enabling the enzymatic reaction to occur *in situ*. Additionally, the limited release of PDH minimizes the risk of unwanted metabolic effects in surrounding healthy tissues while still enabling localized biofilm disruption or metabolic modulation at the wound site.

**Figure 12.**
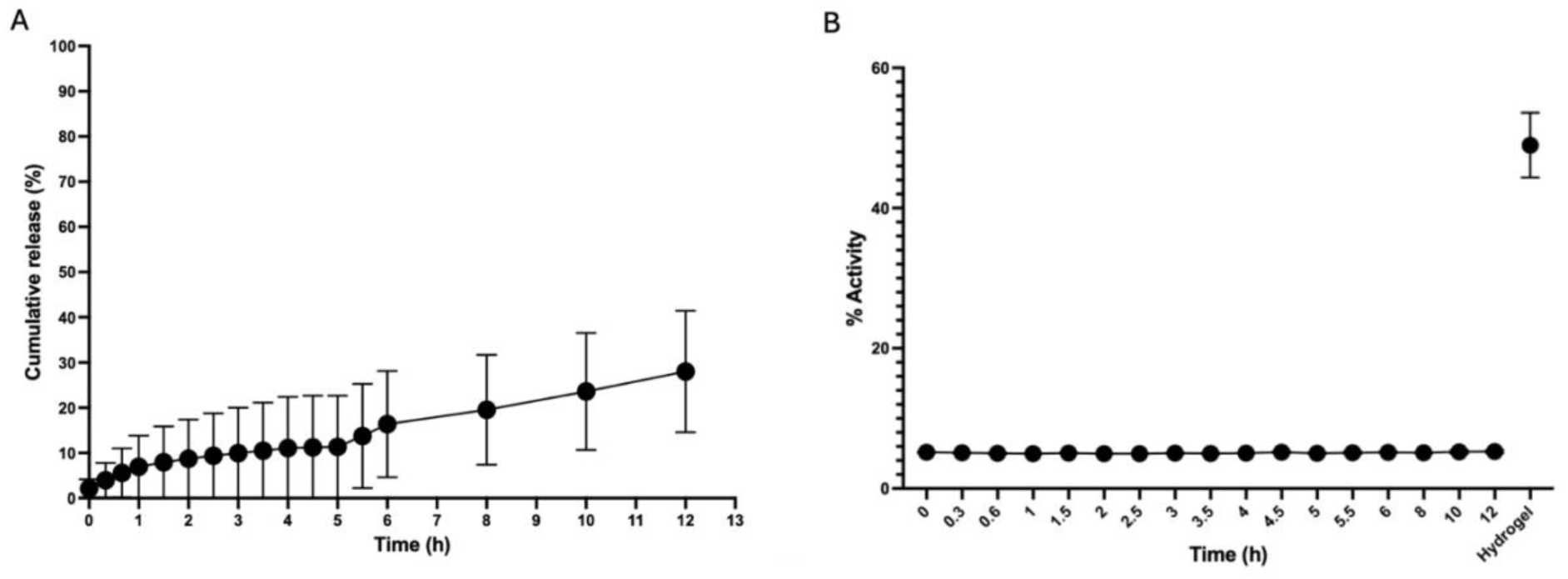
A) Cumulative release percentage of PDH over 12 hours. B) The percentage of enzyme activity compared to initial activity resulting from the released enzyme over 12 hours as well as activity from the hydrogel itself after 12 hours in solution. Data shown are mean ± standard error of the mean. Hydrogels were made in triplicate and Samples taken in duplicate from each hydrogel.

### Alginate Hydrogels Loaded with 0.85 U PDH Significantly Reduce *P. aeruginosa* Biofilms

One of the major pathogens of concern in wounds is *P. aeruginosa*, a Gram-negative bacterium that can establish a biofilm wound infection reaching approximately 1 × 10⁹ CFU per gram of burn tissue within 7 days, regardless of the initial inoculum size [66]. In this biofilm study, alginate hydrogels containing varying concentrations of PDH (0.35 to 0.85 U) were exposed to biofilms, and the biomass after treatment was compared with a commercially available alginate wound dressing (Dimora®) and controls. This concentration range was selected based on the enzyme activity studies shown in Figure 10, which showed that loading less than 0.35 U of PDH resulted in negligible enzymatic activity, while activity was maximized at a loading of 0.85 U.

Among the tested samples, only the hydrogel containing 0.85 U of PDH significantly reduced biofilm biomass, with a decrease of over 40.94 ± 6.54% compared to the control. The unencapsulated PDH sample (5 mU) led to an even greater reduction in biomass. However, the entrapped PDH system has the potential to perform comparably, if not better, under optimized conditions. One limitation of this study was the small volume of hydrogel used for biofilm testing. As shown in Figure 13, using larger volumes of alginate allowed for more enzyme to be entrapped, leading to higher retained enzymatic activity. However, due to the 96-well plate format and the use of peg lids to grow the biofilms, hydrogel volume was restricted to 65 µl to avoid direct contact with the cultured biofilms.

**Figure 13.**
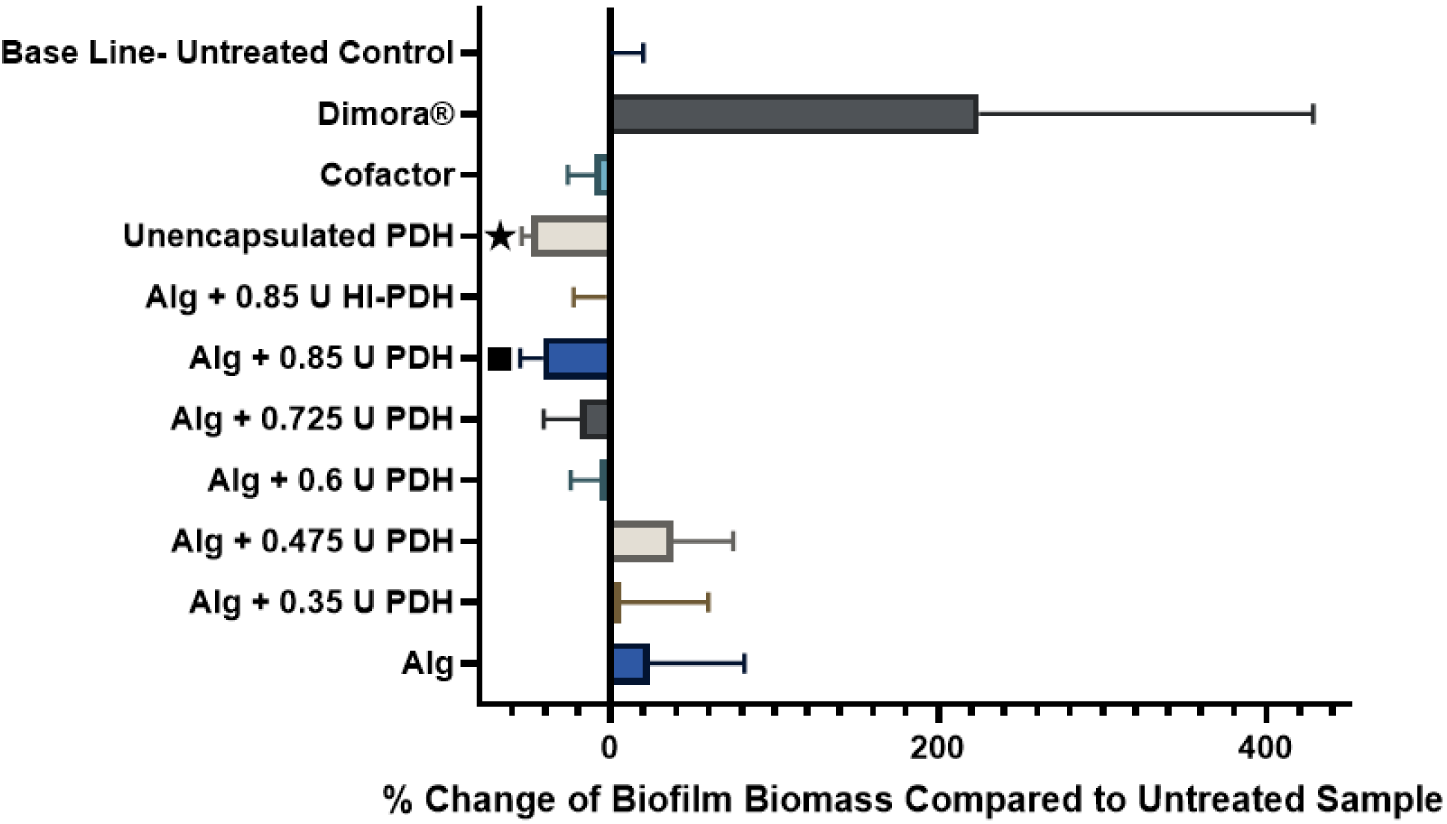
Percentage change of biofilm biomass compared to untreated samples after treatment with PDH-loaded alginate hydrogels containing amounts of 0.35, 0.475, 0.6, 0.725, and 0.85 U PDH as well as 0.85 U of HI-PDH, Dimora® wound dressing, and untreated samples after 24 hours, Data shown are mean ± standard error of the mean. Hydrogels were prepared in triplicate (n=3), Statistically significant results compared to untreated control designated with ▪p < 0.001, *p< 0.0001.

Despite this limitation, a dose-response relationship was observed, with increasing PDH concentrations correlating with greater biofilm biomass reduction. Notably, the alginate-PDH formulation at 0.85 U showed significantly better anti-biofilm performance than the commercial wound dressing, decreasing the biofilm biomass by 80.05 ± 6.37 % comparatively. These results highlight the promise of this technology for biofilm control and support further optimization of formulation volume and enzyme loading to enhance therapeutic potential. Although not tested here, PDH-loaded alginate hydrogels also hold promise to prevent biofilms [26], further highlighting their potential impact.

### Safe and Effective: PDH-Loaded Hydrogels Reduce Biofilms Without Inducing Human Fibroblast Cytotoxicity

To assess the cytotoxicity of the synthesized alginate hydrogels loaded with PDH, confluent HDFa fibroblasts were exposed to leachate dilutions obtained from alginate hydrogels containing 0.85 U of PDH over 24 hours. Cell viability was compared to that of a commercial alginate wound dressing, unloaded alginate hydrogels, and unencapsulated PDH (5 mU). The concentration of 0.85 U was chosen based on previous biofilm and activity studies, which demonstrated maximal enzymatic activity and biofilm biomass reduction at this level. As shown in Figure 14**E****rror! Reference source not found.**, the percent cell viability for the unencapsulated PDH at ½ dilution, PDH-loaded hydrogel, and commercial hydrogel were 73.62 ± 9.10%, 81.52 ± 8.53%, and 88.64 ± 11.98%, respectively. These were the only samples statistically different from the media-only control. These results indicate that while unencapsulated PDH approaches the cytotoxicity threshold of 70% (ISO 10993, [43]), its entrapment in alginate alleviates the toxicity concern. Overall, the data indicate that this hydrogel system not only significantly reduces biofilm biomass but is non-cytotoxic to human dermal fibroblasts.

**Figure 14.**
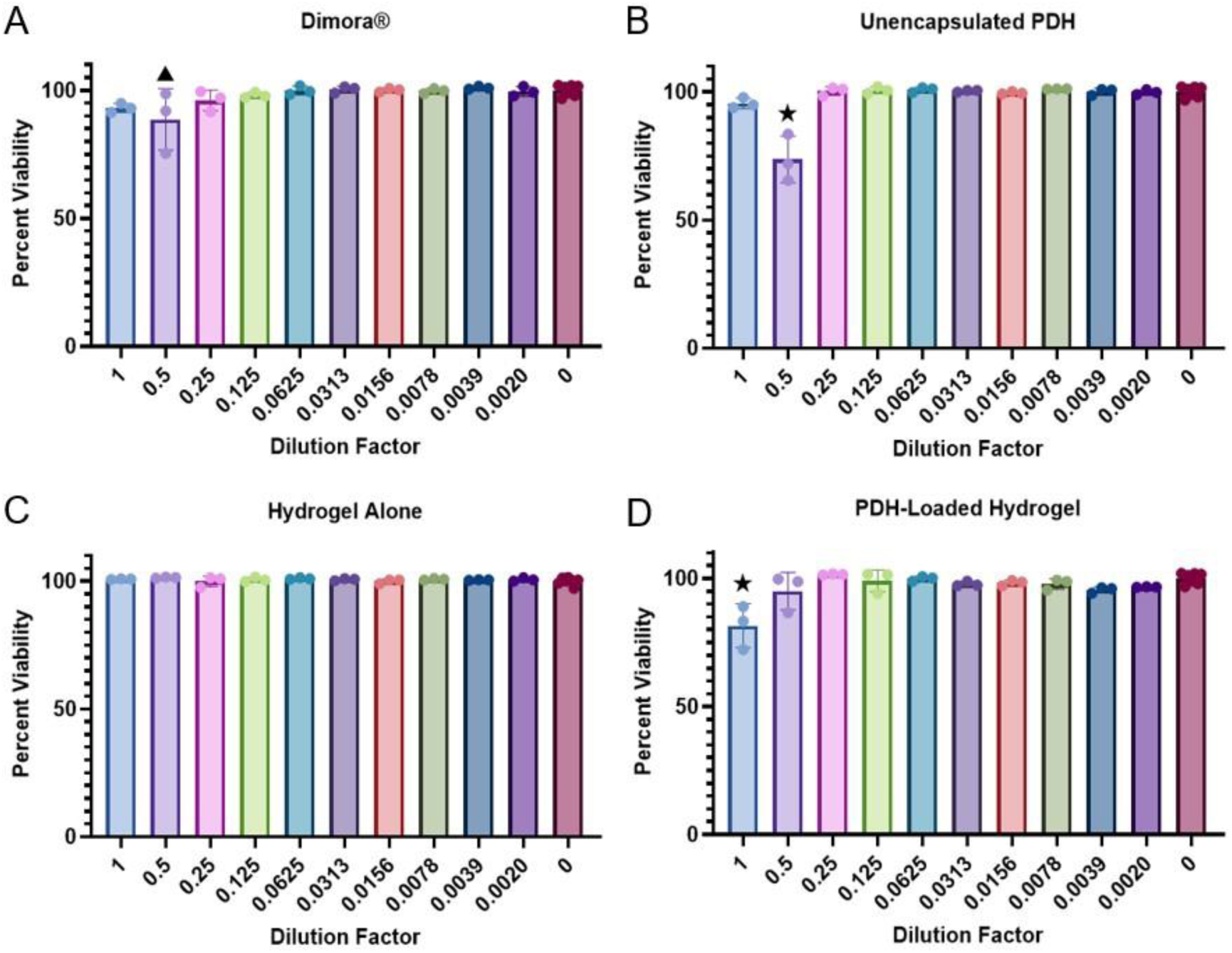
The percent viability compared to cell-only control of HDFa cells exposed to leachate dilutions for 24 hours of A) Dimora® commercial wound dressing, B) unencapsulated PDH, C) alginate hydrogels prepared without encapsulant, and D) PDH-loaded hydrogel. Data shown are mean ± standard error. Leachate samples were taken in triplicate from individual hydrogels (n=3). Statistical significance denoted as pp<0.01 and *p<0.0001 compared to 0 dilution factor.

## Conclusion

This study demonstrates the feasibility of using PDH-loaded alginate hydrogels to disrupt bacterial biofilms, offering a novel strategy for chronic wound management. The optimized hydrogel formulation not only met the mechanical and swelling criteria for wound applications but also successfully retained enzymatic activity over several days. The significant reduction in *P. aeruginosa* biofilm biomass, particularly in comparison to commercial alginate dressings, underscores the therapeutic potential of this metabolism-altering approach. Moreover, the improved biocompatibility observed with encapsulated PDH supports its suitability for direct wound contact. However, a notable limitation of this system is its relatively low encapsulation efficiency, which may limit the amount of active enzyme delivered per unit volume. Future work should focus on improving encapsulation strategies—potentially through surface modification, co-polymers, or microstructural optimization—to enhance loading efficiency without compromising release kinetics or bioactivity. Additionally, *in vivo* validation and studies investigating synergistic effects with antibiotics will be essential to fully realize the clinical potential of this platform. Collectively, this work advances the field of biofilm-interfering wound dressings by establishing a hydrogel system capable of sustained enzyme delivery and targeted biofilm disruption.

## Acknowledgments

This work was supported by the National Science Foundation CAREER Award 2341706. The authors thank Drs. Matthew White, Linda S. Schadler, and Rachael Floreani for the use of the SEM, DHR-2 rheometer, and AR2000 rheometer, respectively. We would like to thank Dr. Matthew J. Wargo for generously providing the PAO1 for this study. The authors also thank Patrick Charron for the training he provided on both rheometer instruments.

